# Auditory and visual object processing in olfactory cortex of individuals with life-long olfactory deprivation

**DOI:** 10.64898/2025.12.16.694602

**Authors:** Evelina Thunell, Moa G. Peter, Fahimeh Darki, Johan N. Lundström

## Abstract

The traditional view of modality-specific brain organization has been challenged by demonstrations of cross-modal processing, i.e. the activation of sensory cortex by atypical sensory input. An alternative theory suggests that rather than specializing simply in different sensory modalities, different parts of the cortex can specialize in different types of tasks. We recently addressed this question in the olfactory domain, showing that in normosmic individuals, the primary olfactory cortex (piriform cortex) responds to unimodal auditory and visual objects, regardless of how strongly they are associated with an odor. Here, we assess whether previous olfactory experience is a prerequisite for this cross-modal processing in the piriform cortex, using a unique group of individuals born without a sense of smell (congenital anosmia; CA; *n* = 30). First, we were able to confirm the presence of clear visual and auditory activations in the piriform cortex of these sensory deprived individuals. As compared to normosmic controls (*n* = 30), these activations and associated functional connectivity, both within the piriform cortices and between piriform cortex and other regions, were altered in a modality dependent way. Our results show that life-long absence of olfactory input does not impede cross-modal activations by visual and auditory objects in the piriform cortex and are compatible with the idea of a task-based rather than modality-specific organization of this brain region.

## 1. INTRODUCTION

Our sensory cortices have traditionally been viewed as predominantly modality-specific, i.e., dedicated to processing input from their respective sensory modalities. However, accumulating evidence from both single-cell level and human neuroimaging studies challenges this view, showing that the sensory cortices also respond to atypical stimuli, e.g. the visual cortex responding to auditory stimuli; a phenomenon known as cross-modal processing (Teichert & Bolz, 2018). A commonly proposed ecological function of cross-modal processing in individuals with intact senses is to facilitate the processing of typical sensory information by evoking associations or imagery of the typical stimuli, or by influencing the processing depending on the congruency of the typical and atypical stimuli. An alternative viewpoint that has emerged is that the sensory cortices specialize in different types of tasks rather than different sensory modalities (Amedi et al., 2017). For example, the visual cortex demonstrates excellent spatial processing abilities whereas the auditory cortex is better suited for temporal processing. Studies on individuals with congenital blindness and deafness suggest that the deprived sensory cortex sustains the same type of tasks as in sighted and hearing individuals but based on atypical sensory stimuli (Amedi et al., 2017): The processing related to localization, tools, and objects in higher-order “visual” cortex remains remarkably consistent in congenitally blind individuals even when the sensory information stems from non-visual modalities (Cecchetti et al., 2016; Gougoux et al., 2005; Heimler et al., 2015; Renier et al., 2010; Weeks et al., 2000).

Analogously, in deaf individuals, activation of the auditory cortex by visual motion has been demonstrated (Fine et al., 2005), and (Bola et al., 2017) found that rhythmic visual input activated the auditory cortex in a similar way to rhythmic auditory input in hearing controls.

We recently demonstrated cross-modal processing in the olfactory cortex in normosmic individuals (Thunell et al., 2025). Specifically, the posterior piriform cortex (pPir), a region associated with olfactory object and category processing (Gottfried, 2010; Howard et al., 2009) was activated by both unimodal auditory and unimodal visual object stimuli, and the anterior piriform cortex (aPir) was activated by auditory stimuli. In earlier reports of piriform activation by non-olfactory stimuli, the stimulation had some type of olfactory significance (González et al., 2006; Gottfried et al., 2004; Porada et al., 2019; Schulze et al., 2017). For example, (Kehl et al., 2024) demonstrated that the human piriform cortex codes not only for specific odors, but also for pictures and written names of objects corresponding to those odors. Our recent study (Thunell et al., 2025) shows that cross-modal processing in the piriform cortex can exist even in the absence of olfactory context and, interestingly, the level of activation did not depend on the odor association of the presented stimuli, indicating that it is not merely a reflection of evoked odor imagery. Still, it could be a result of a lifetime of exposure to non-olfactory stimuli with olfactory significance: Repeated coactivation of olfactory and other sensory cortices may have strengthened their overall connections and the cross-modal activity can therefore be evoked even by stimuli with very minor olfactory association. This raises the question: Is the sense of olfaction a prerequisite for cross-modal processing in the piriform cortex?

Here, we use a unique population of individuals with isolated congenital anosmia (CA; *n* = 30) and matched normosmic controls (*n* = 30) to assess whether non-olfactory processing in the piriform cortex requires experience of olfactory input or whether it is an inherent feature. If the non-olfactory processing in the piriform cortex in normosmics is a result of repeated olfactory-multisensory input, we expect to be strongly reduced or absent in individuals with CA compared to normosmic controls.

## 2. MATERIALS AND METHODS

A subset of the fMRI data from the control group were previously used in (Thunell et al., 2025). Data are available on the OSF repository: https://osf.io/4t6y5/overview

### 2.1 Participants

A total of 60 participants were included in this study: 30 otherwise healthy individuals with isolated congenital anosmia (CA, congenital anosmia unrelated to specific genetic disorders such as Kallmann syndrome; convenience sample) and 30 healthy controls, matched in terms of age and gender (Table 1). Inclusion criteria for the CA group were a self-reported lifelong lack of olfactory perception (no memories of ever smelling an odor) without any known underlying condition causing the anosmia, such as medical issues or experiences associated with a risk of olfactory deterioration, as ascertained by a structured interview, and scoring in the anosmia range in the olfactory behavioral test (see Olfactory screening below). The inclusion criterion for the Control group was a self-proclaimed functional sense of smell and scoring in the normosmia range in the olfactory behavioral test. In addition, all participants in both groups reported having normal or corrected-to normal visual acuity and normal hearing. The study was approved by the Swedish Ethical Review Authority [Dnr. 2020-06533] and performed in accordance with the Helsinki declaration. All participants provided written informed consent prior to the experiment.

**Table 1.** Descriptive statistics per experimental group.

|  | CA (n = 30) | Control (n = 30) |
| --- | --- | --- |
| Age (years) | 35.8 ± 9.7 | 35.4 ± 10 |
| Women | 17 | 17 |
| Men | 13 | 13 |
| TDI | 9.5 ± 2.4 | 35.9 ± 3.3 |
| *Threshold* | 1.2 ± 0.5 | 9.4 ± 2.3 |
| *Discrimination* | 4.3 ± 1.5 | 12.8 ± 2 |
| *Identification* | 4 ± 1.7 | 13.7 ± 1.1 |

Values presented as mean ± standard deviation. CA = congenital anosmia, TDI = combined score from the three Sniffin' Sticks olfactory subtests (Threshold, Discrimination, and Identification).

### 2.2 Experimental Design

#### 2.2.1 Stimuli

The stimulus set consisted of matching visual (grayscale pictures) and auditory (sound clips) stimuli representing 48 common and, based on a pilot study on 23 healthy participants, easily identifiable objects (for details, see (Thunell et al., 2025)). Pictures were obtained from the Open Image Dataset (https://opensource.google) and ImageNet (http://www.image-net.org) and symmetrically cropped to a square shape with the shortest of the original height and width defining the resulting side lengths. All images were converted to grayscale using the rgb2gray function in Matlab to eliminate color variation. Sounds were obtained from either the BBC Sounds Database (http://bbcsfx.acropolis.org.uk) or Freesound (https://freesound.org) and either trimmed or seamlessly looped to last 3 seconds. They were subsequently equalized with a root mean square equalization of -23 dB loudness (Regenbogen et al. 2016). In the magnetic resonance (MR) scanner, pictures were presented on a 40" 4K ultra-HD LCD display (NordicNeuroLab AS, Bergen, Norway) placed behind the scanner bore and viewed via a head coil mirror and spanned approximately 4.5 degrees of visual angle. Sounds were presented via binaural headphones (NordicNeuroLab AS, Bergen, Norway).

#### 2.2.2. Task

To ensure that all presented objects were familiar, participants were presented with all stimuli one at a time together with the name of the object in a practice session prior to MR scanning. To maximize object recognition, they were instructed to learn to identify the objects and incorrectly told that they would later be tested on this task.

The functional part of the MR scanning session included four runs, each lasting 10 min and 15 s and containing eight blocks, each lasting 1 min. Each block was both preceded and followed by 15 s of no stimulus presentation. A block consisted of six stimulus presentations and included only auditory or only visual stimuli. Stimulus duration was 3 s and each stimulus presentation was preceded by 7 s ± 3 s of no stimulus. All stimuli (48 sounds and 48 pictures) were presented twice; once during the first half of the runs and once during the second half of the runs, in a random order, meaning that each participant was exposed to a total of 192 events (96 auditory and 96 visual stimulus presentations). Participants were instructed to fixate their gaze on a central cross which was visible throughout each run except during picture presentation, and to silently identify each presented object without providing any response, to limit motor-related effects. All participants completed all four functional runs, except for one CA participant who completed only three.

#### 2.2.3 Olfactory screening

After the MR scanning, participants’ olfactory functions were assessed with the Sniffin’ Sticks olfactory test (Burghart, Wedel, Germany) to ensure functional anosmia in the CA group (all classified as anosmic) and a normal sense of smell in the Control group (all classified as normosmic in their respective age group; Table 1). The Sniffin’ Sticks test is a clinically validated test that consists of three subtests with individual scores: odor detection threshold (T), odor quality discrimination (D), and 4-alternative cued odor quality identification (I). The total TDI score is the sum of these three scores and is used to classify individuals as normosmic, hyposmic, or anosmic based on normative data from over 9000 individuals (Oleszkiewicz et al., 2019). Individuals with CA who had already gone through the full Sniffin’ Sticks olfactory test as part of one of our previous studies were exempt from olfactory screening and their previous score was instead used.

### 2.3 MR imaging

Imaging data were acquired using a 3T Siemens Magnetom Prisma MR scanner (Siemens Healthcare, Erlangen, Germany) with a 20-channel head coil. Functional data were acquired using 3D echo planar sequences with the following settings: 56 slices, TR = 1700 ms, TE = 30 ms, FA = 70°, voxel size = 2.234 mm x 2.234 mm x 2.200 mm, and FoV = 94 x 94 voxels. Structural T_1_-weighted images were acquired with whole-brain coverage using a 3D GR/IR T1-weighted sequence with 208 slices, TR = 2300 ms, TE = 2.89 ms, FA = 9°, voxel size = 1 mm x 1 mm x 1 mm, and FoV = 256 x 256 voxels.

### 2.4 MR image analysis

#### 2.4.1 Preprocessing

Preprocessing of the MR data was done using SPM12 (ver. 7487, Wellcome Trust Centre for Neuroimaging, UCL; https://www.fil.ion.ucl.ac.uk/spm/) in Matlab 2019b (MathWorks, Natick, Massachusetts, USA) and included slice timing correction to address differences in timing of slice acquisition in each functional volume, realignment of functional volumes within each run to account for movement, coregistration of the functional runs, coregistration of the anatomical to the functional images, and normalization to MNI space. Finally, the functional images were smoothed with a 6 mm full width at half maximum (FWHM) Gaussian kernel.

#### 2.4.2 Region of interest and exploratory whole-brain analyses

To assess cross-modal processing in olfactory cortex after lifelong complete olfactory sensory deprivation, and to test whether it is altered as compared to controls, we used a region of interest (ROI) approach targeting the anterior piriform cortex (aPir), known to code for basic odor properties, and the posterior piriform cortex (pPir), which processes odor objects (Gottfried, 2010; Howard et al., 2009; Lundström et al., 2011), as ROIs (Figure 1A). To ensure that our experimental paradigm and data processing worked as intended, and to assess whether potential group differences were unique to the olfactory sensory system, we also included two auditory and two visual ROIs. The auditory and visual ROIs were chosen to represent similar functions as the olfactory regions: Representation of basic stimulus properties (i.e., aPir) in the primary auditory (A1) and primary visual (V1) cortex, and higher order processing and object representation (i.e., pPir) in the higher-order auditory cortices (hAC) and the lateral occipital complex (LOC; Figure 2A, D). Definition and creation of the six ROIs are described in detail in (Porada et al., 2019). Finally, to explore effects outside of our olfactory and control ROIs, we conducted exploratory whole-brain voxel-wise group comparisons.

**Figure 1:**
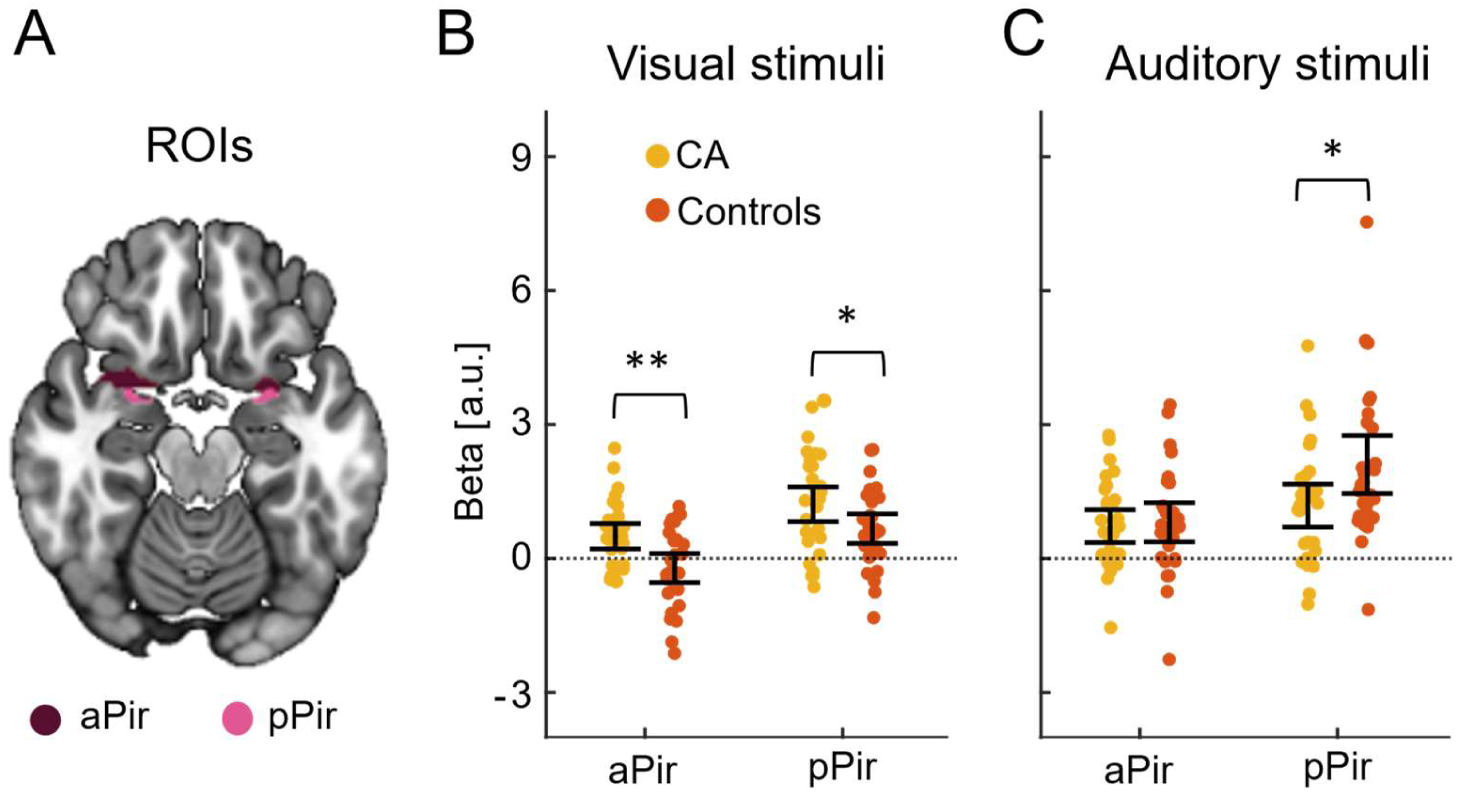
Visual and auditory processing in olfactory (piriform) cortex. **A)** Locations of the anterior and posterior piriform cortex (aPir and pPir) regions of interest. **B)** Estimated beta values for visual activation in aPir and pPir for each individual with Congenital Anosmia (CA) and in the Control group. Error bars indicate 95% confidence intervals. **C)** Estimated beta values for auditory activation in aPir and pPir. Error bars denote 95% confidence intervals. Group comparisons: *p < .05, **p < .01.

**Figure 2:**
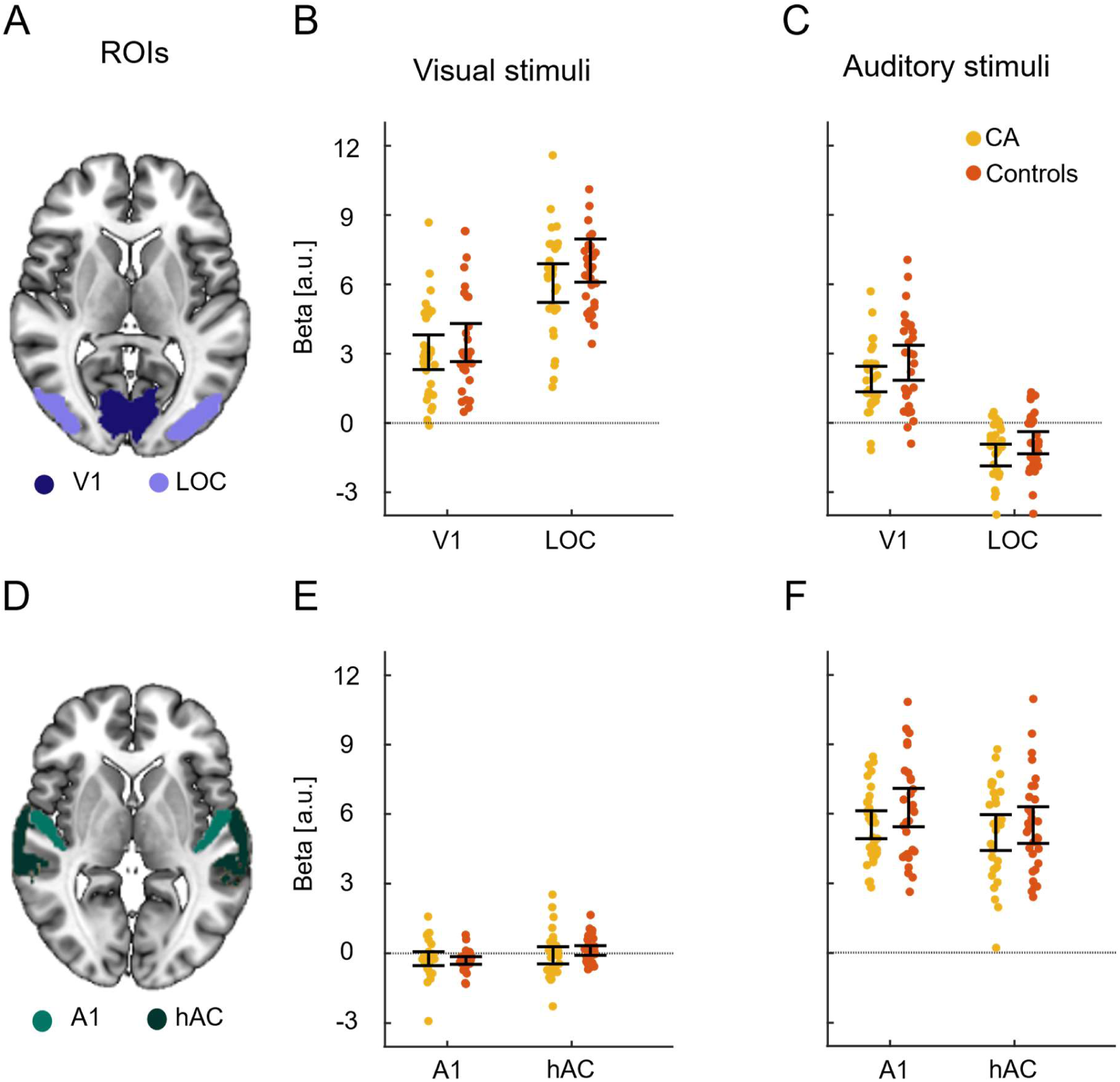
Visual and auditory processing in visual and auditory cortices. **A)** Locations of primary visual cortex (V1) and lateral occipital complex (LOC) regions of interest. **B)** Estimated beta values for visual processing in V1 and LOC for each individual with congenital anosmia (CA) and in the Control group. **C)** Estimated beta values for auditory processing in V1 and LOC. **D)** Locations of primary auditory cortex (A1) and higher-order auditory cortices (hAC) regions of interest. **E)** Estimated beta values for visual processing in A1 and hAC. **F)** Estimated beta values for auditory processing in A1 and hAC. Error bars indicate 95% confidence intervals.

Activity related to auditory and visual processing was modelled for each participant using a general linear model (GLM) implemented in SPM12. The model included two conditions, Visual and Auditory stimulation, modelled as 3 s long boxcar functions convolved with a canonical hemodynamic response function, and the six realignment parameters derived from the preprocessing realignment step as nuisance regressors to account for effects of participant motion. A high-pass filter (cutoff 128 s) was added to remove low frequency drift, and a first-order autoregressive model was used to account for serial correlations. Auditory and visual activations were calculated voxel-wise for each participant against the implicit baseline. These analysis steps were conducted for both non-smoothed and smoothed images for subsequent ROI and whole-brain analysis, respectively.

For each participant, mean contrast estimates (beta values) were extracted for unilateral ROIs and then averaged over the left and right hemispheres for the two conditions (Visual and Auditory stimulation). To test our hypothesis of group differences in auditory and visual cross-modal processing in the piriform cortex, mean betas in aPir and pPir were compared using Welch’s *t*-tests with an α criterion of .05. Control analyses of potential group differences in auditory and visual processing in our auditory and visual ROIs were conducted following the same steps as for the olfactory ROIs. In the exploratory whole-brain voxel-wise group comparisons, we used a family-wise error (FWE) corrected threshold of *p* < .05.

#### 2.4.3 Functional connectivity analysis

To assess how olfactory regions communicate with other brain regions during cross-modal processing in individuals with life-long absence of olfactory input, we conducted both ROI-to-ROI and ROI-to-whole-brain functional connectivity analyses using the CONN toolbox (Whitfield-Gabrieli & Nieto-Castanon, 2012). For the ROI-to-ROI analysis, we considered both the olfactory ROIs (aPir and pPir) and control regions (V1, LOC, A1, and hAC). A weighted general linear model (wGLM) was used to estimate pairwise ROI-to-ROI connectivity separately for each condition (Visual and Auditory) by computing bivariate correlations across all time-points restricted to the duration of the stimulation. White matter and cerebrospinal fluid time series as well as head movements were regressed out for each participant to account for physiological noise. Following the calculation of the functional connectivity between all ROIs across the two conditions, we tested for group differences using t-tests (α criterion = .05). All second-level group comparison results were corrected for multiple comparisons using the false discovery rate (FDR) procedure (Benjamini & Hochberg, 1995) with a threshold of q <.05.

We next conducted a ROI-to-whole-brain analysis to assess the functional connectivity of aPir and pPir to the whole brain, using a weighted seed-based connectivity (wSBC) approach to account for condition-specific functional connectivity. The mean time-series of each ROI was correlated with the time-series of all voxels across the entire brain, restricted to the duration of each condition (Visual and Auditory). A whole-brain voxel-wise group comparison was then performed using a t-test for the two conditions, at the *p* < .05 family-wise error (FWE) level.

## 3. RESULTS

### Altered cross-modal processing in olfactory cortex in congenital anosmia

First, we assessed whether non-olfactory processing in olfactory (piriform) cortex, which has previously been demonstrated in normosmic individuals, is present also in individuals with lifelong olfactory deprivation. Indeed, we found strong cross-modal activations in both the anterior piriform cortex (aPir) and posterior piriform cortex (pPir) in both the normosmic Control and Congenital Anosmia (CA) groups: The auditory and visual stimulation elicited significant activations in both aPir and pPir in both groups (all *p’s* < .002, Supplementary Table S1), with the exception of visual processing in aPir in the Control group (*p* = .186). Next, we compared the activations between the groups: We found significantly higher visual activation in the CA group compared to the Control group in both aPir, *t*(56.9) = 3.38, *p* = .001, *d* = 0.87, and pPir, *t*(56.6) = 2.18, *p* = .033, *d* = 0.56 (Figure 1B). For the auditory stimulation, the activation in the pPir was lower in the CA group *t*(53.4) = 2.33, *p* = .024, *d* = 0.6 (Figure 1C), whereas there was no significant group difference in aPir, *t*(56.4) = 0.3, *p* = .763, *d* = 0.08 (Figure 1C).

To assess whether the altered cross-modal processing in individuals with CA is limited to the olfactory regions, we also analyzed the visual and auditory ROIs. We did not find any significant group differences for either auditory activation in the visual ROIs, or visual activation in the auditory ROIs (Figure 2, Table 2). Additional Bayesian t-tests of the crossmodal effects show moderate evidence for the lack of visual processing in primary auditory cortex (A1; B_01_ = 3.47), and anecdotal evidence for the remaining null effects (visual processing in higher auditory cortex (hAC): B_01_ = 2.51, auditory processing in primary visual cortex (V1): B_01_ = 1.43, and auditory processing in the lateral occipital cortex (LOC): B_01_ = 1.27). In addition, we did not observe any significant group differences in the typical activations (auditory activation in the auditory ROIs and visual activation in the visual ROIs; Figure 2, Table 2). The exploratory whole-brain voxel-wise analysis did not reveal any group differences for either auditory or visual processing.

**Table 2.** Mean beta results from ROI analysis, per condition, ROI partition, and participant group. Statistically significant group differences are denoted by *.

| Region of Interest | Mean beta ± SD | | df | *t* | *p* | *d* |
| --- | --- | --- | --- | --- | --- | --- |
|  | Congenital Anosmia | Control |  |  |  |  |
| *Visual stimuli* |  |  |  |  |  |  |
| *aPir* | 0.5 ± 0.76 | -0.22 ± 0.87 | 56.9 | 3.38 | .001* | .87 |
| *pPir* | 1.21 ± 1.04 | 0.67 ± 0.88 | 56.6 | 2.18 | .033* | .56 |
| *V1* | 3.06 ± 2.01 | 3.48 ± 2.2 | 57.5 | 0.77 | .442 | .20 |
| *LOC* | 6.06 ± 2.25 | 7.03 ± 2.5 | 57.3 | 1.59 | .118 | .41 |
| *A1* | -0.21 ± 0.80 | -0.29 ± 0.44 | 45.2 | 0.47 | .641 | .12 |
| *hAC* | -0.07 ± 1.00 | 0.14 ± 0.56 | 45.6 | 1.00 | .324 | .26 |
| *Auditory stimuli* |  |  |  |  |  |  |
| *aPir* | 0.73 ± 0.99 | 0.81 ± 1.17 | 56.4 | 0.30 | .763 | .08 |
| *pPir* | 1.19 ± 1.29 | 2.11 ± 1.74 | 53.4 | 2.33 | .024* | .60 |
| *V1* | 1.90 ± 1.48 | 2.6 ± 2.02 | 53.1 | 1.53 | .131 | .40 |
| *LOC* | -1.39 ± 1.26 | -0.86 ± 1.28 | 58 | 1.62 | .110 | .42 |
| *A1* | 5.52 ± 1.61 | 6.26 ± 2.22 | 53 | 1.48 | .145 | .38 |
| *hAC* | 5.18 ± 2.06 | 5.51 ± 2.11 | 58 | 0.60 | .550 | .16 |

To determine whether the observed differences in cross-modal piriform activation between CA and Control individuals reflect broader network-level changes, we assessed the functional connectivity between the six regions of interest (aPir, pPir, V1, LOC, A1, and hAC). We found stronger connectivity between aPir and pPir in the CA group compared to Controls, during both visual and auditory stimulation (Fig. 3, Table 3). The connectivity between V1 and both piriform ROIs was instead decreased in the CA individuals, for both auditory and visual stimulation. Between LOC and pPir, we found weaker connectivity in the CA group than in the Controls during visual stimulation, but no group difference during auditory stimulation. There were no significant group differences in connectivity between the two visual areas (V1 and LOC). The connectivity between the two auditory ROIs (A1 and hAC) was higher in CA compared to Controls during auditory stimulation, whereas there was no significant group difference during visual stimulation.

**Figure 3:**
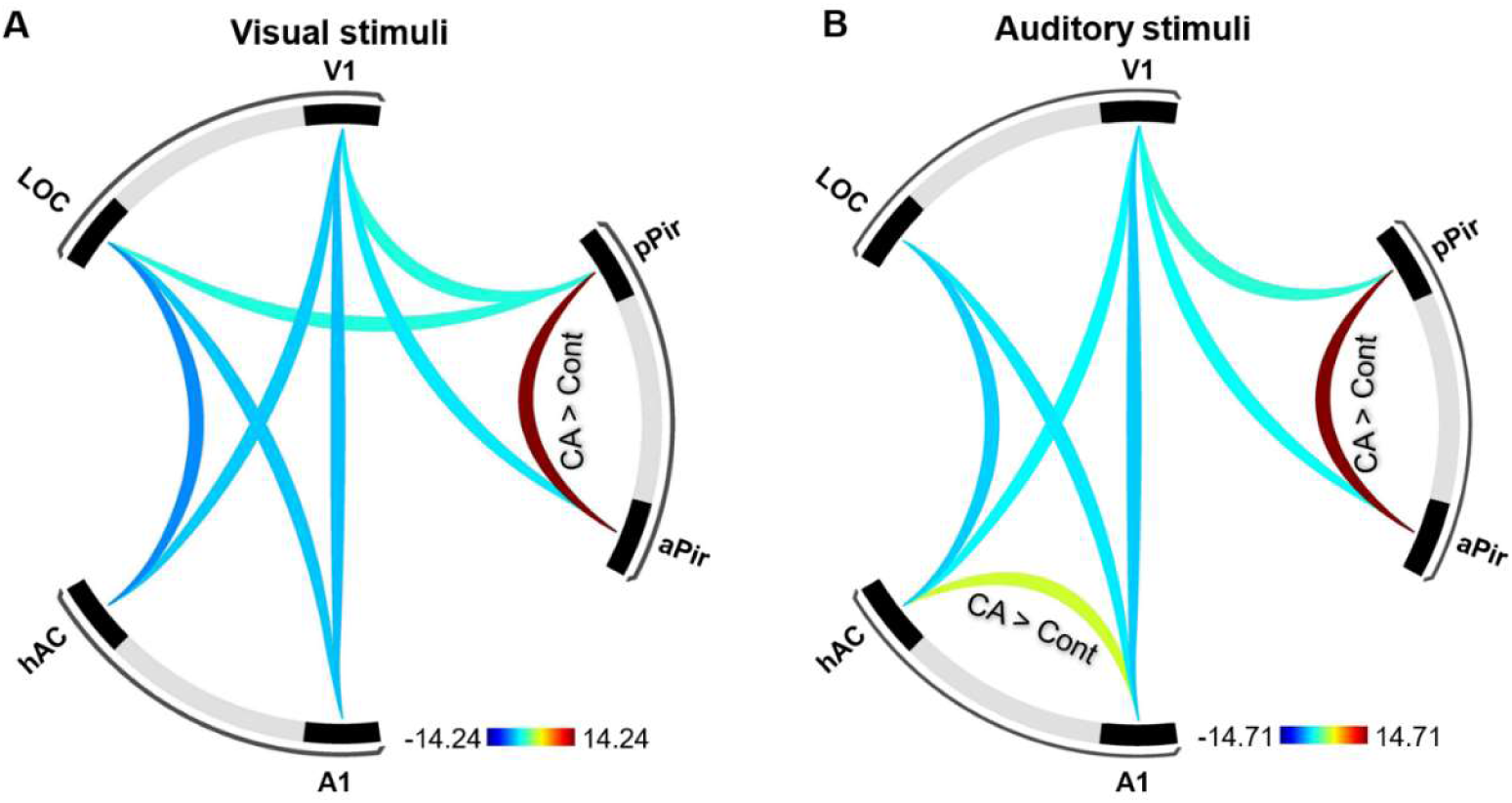
Statistically significant group differences in ROI-to-ROI functional connectivity. Connections that are stronger in the congenital anosmia (CA) group compared to Controls (Cont) are marked CA > Cont and drawn in warm colors; connections that are stronger in controls than in the congenital anosmia group are drawn in colder colors. The color scale depicts T values.

**Table 3.** Group comparisons of ROI-to-ROI functional connectivity. The table reports t-statistics (T), uncorrected p-values (p-unc), and FDR-corrected p-values (5%, p-FDR) for pairwise ROI-to-ROI connectivity comparisons between the congenital anosmia (CA) group and controls (Cont.), during both visual (left) and auditory stimulation (right).

| ROI-ROI | Visual stimuli |  |  |  | Auditory stimuli |  |  |  |
| --- | --- | --- | --- | --- | --- | --- | --- | --- |
|  | T | difference | p-unc | p-FDR | T | difference | p-unc | p-FDR |
| aPir - pPir | 14.24 | CA > Cont. | $< 10^{-6}$ | $< 10^{-6}$ | 14.71 | CA > Cont. | $< 10^{-6}$ | $< 10^{-6}$ |
| hAC - LOC | -6.75 | Cont. > CA | $< 10^{-6}$ | $< 10^{-6}$ | -5.19 | Cont. > CA | $3.00 \times 10^{-6}$ | $1.50 \times 10^{-5}$ |
| A1 - V1 | -5.32 | Cont. > CA | $2.00 \times 10^{-6}$ | $8.00 \times 10^{-6}$ | -5.09 | Cont. > CA | $4.00 \times 10^{-6}$ | $2.10 \times 10^{-5}$ |
| A1 - LOC | -5.17 | Cont. > CA | $3.00 \times 10^{-6}$ | $8.00 \times 10^{-6}$ | -4.25 | Cont. > CA | $8.10 \times 10^{-5}$ | $2.03 \times 10^{-4}$ |
| hAC - V1 | -5.02 | Cont. > CA | $5.00 \times 10^{-6}$ | $1.30 \times 10^{-5}$ | -3.81 | Cont. > CA | $3.39 \times 10^{-4}$ | $8.47 \times 10^{-4}$ |
| aPir - V1 | -4.10 | Cont. > CA | $1.35 \times 10^{-4}$ | $3.37 \times 10^{-4}$ | -3.47 | Cont. > CA | $1.01 \times 10^{-3}$ | $2.53 \times 10^{-3}$ |
| pPir - V1 | -2.66 | Cont. > CA | .01 | 0.02 | -2.50 | Cont. > CA | .02 | .04 |
| pPir - LOC | -2.62 | Cont. > CA | .01 | 0.02 | 2.28 | CA > Cont. | .03 | .04 |

To explore whether the altered pattern of connectivity in the CA group extends to other brain regions, we also examined ROI to whole brain functional connectivity (Fig. 4). The functional connection between the olfactory (piriform) cortex (aPir and pPir) and amygdalae was stronger in the CA group than in Controls, during both visual and auditory stimulation. In addition, connectivity between the piriform cortex and occipital/parietal regions was overall reduced in the CA group.

**Figure 4:**
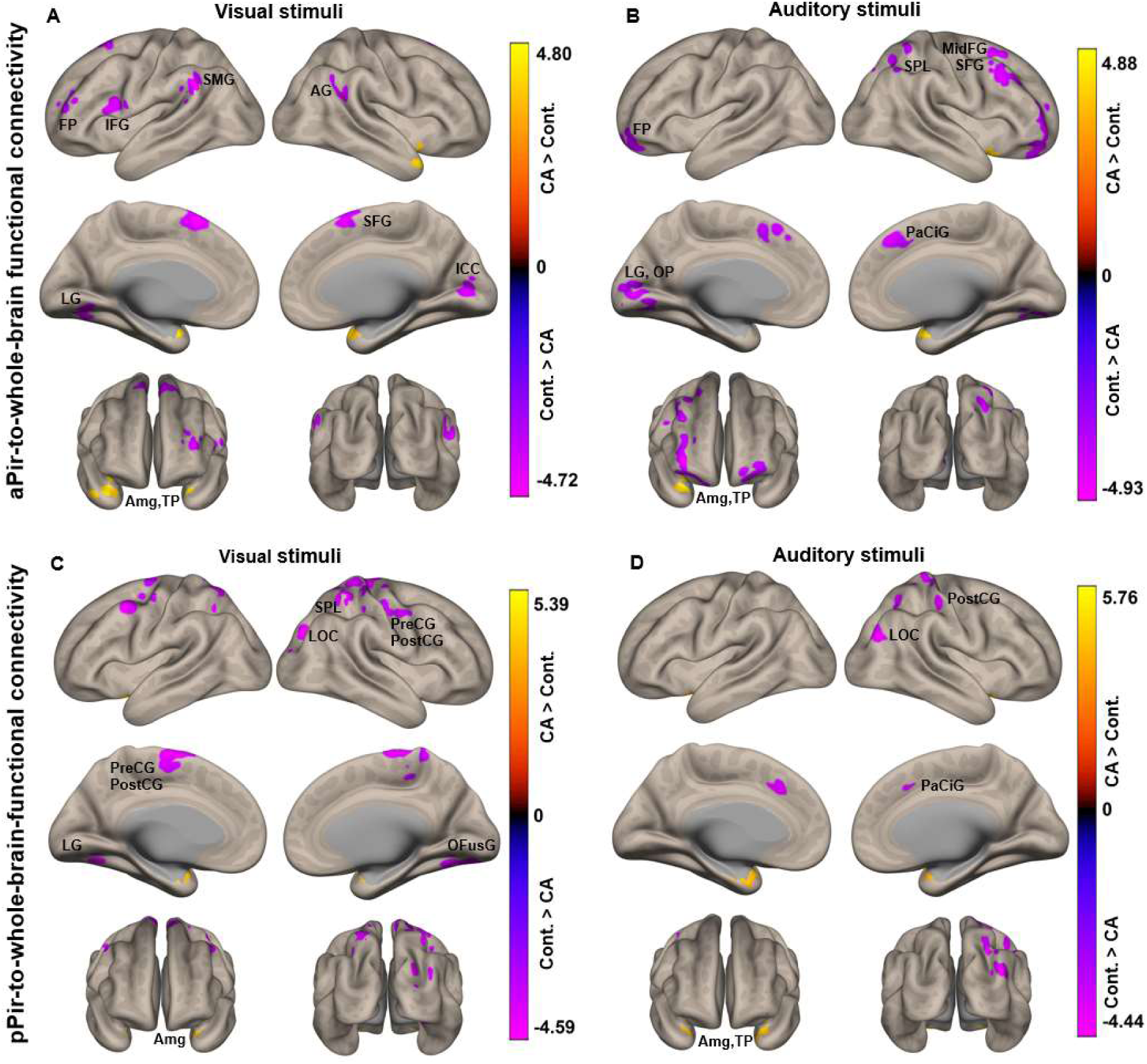
Whole-brain functional connectivity patterns of the anterior and posterior piriform cortex. (A) and (B) show whole-brain functional connectivity maps seeded from the anterior piriform cortex (aPir) during visual and auditory stimulation, respectively. (C) and (D) display whole-brain connectivity to the posterior piriform cortex (pPir). The color scale depicts T values. FP = frontopolar cortex, IFG = inferior frontal gyrus, SMG = supramarginal gyrus, AG = angular gyrus, LG = lingual gyrus, SFG = superior frontal gyrus, ICC = intracalcarine cortex, Amg = amygdala, TP = temporal pole, SPL = superior parietal lobule, LOC = lateral occipital complex, preCG = precentral gyrus, postCG = postcentral gyrus, OfusG = occipital fusiform gyrus, midFG = middle frontal gyrus, OP = occipital pole, PaCiG = paracingulate gyrus.

## 4. DISCUSSION

We here demonstrate activation by unisensory visual and auditory objects in the piriform cortex of individuals born without a sense of smell (congenital anosmia; CA), an effect that has previously only been demonstrated in normosmic individuals (Thunell et al., 2025). This finding demonstrates that the development of visual and auditory cross-modal processing in the piriform cortex does not require any past experience of olfactory stimulation.

These cross-modal activations differed between the CA group and the normosmic control group: In individuals with CA, the visual activation was higher in both the anterior and posterior piriform cortex, while auditory activation was lower in posterior piriform cortex. Thus, the plasticity of the piriform cortex in CA is more complex than a simple overall up- or down-regulation of the cross-modal processing. There were no significant group differences in the visual or auditory ROIs, or in the whole-brain analysis, suggesting that the altered cross-modal processing associated with lifelong absence of olfactory input may be specific to the piriform cortex. Our previous structural analyses suggest that this effect is not attributable to any group differences in piriform morphology (Peter et al., 2020, 2023), but see also (Frasnelli et al., 2013); instead, these results might be related to our findings of altered connectivity within the piriform cortex (between aPir and pPir), and between the piriform cortex and occipital/parietal regions, including the visual ROIs, and the amygdalae. Possibly, the visual modality exerts a takeover effect in the piriform cortex in individuals with CA, motivated by its relatively important object-related information. Indeed, vision has been found to be involved in olfactory object processing, where repetitive transcranial magnetic stimulation (rTMS) of the occipital cortex facilitates the discrimination of odor quality but not intensity (Jadauji et al., 2012). In contrast, rTMS to the auditory cortex did not have any such effect. In rodents, the coupling of beta band oscillations between auditory and olfactory cortex seems to be learned (Olofsson et al., 2019), suggesting that olfactory coactivation may be needed for auditory connections to olfactory cortex to strengthen.

In the visual and auditory systems, the size of the primary cortices has been found to correlate with perception. For example, the surface area of primary visual cortex (V1) has been found to predict subjective experience of object size as well as the strength of some illusions (Moutsiana et al., 2016; Schwarzkopf & Rees, 2013; Schwarzkopf et al., 2011). In the primary auditory cortex, surface area has been found to correlate with pitch perception and general auditory abilities (Palomar-García et al., 2020; Schneider et al., 2005). On the contrary, there are to our knowledge only reported null findings of correlation effects between the structure of the piriform cortex and olfactory performance, suggesting that the olfactory cortex might be less of a unimodal processing area than the auditory and visual cortices. For example, olfactory ability has been shown to correlate with orbitofrontal cortex (OFC) and olfactory bulb volume but not with piriform cortex volume (Seubert et al., 2013; Shen et al., 2013).

A dominant theory of cross-modal plasticity in sensory deprived individuals is that the deprived cortex reorganizes to compensate for the missing sense by allocating more computational power to one or more of the remaining senses. Alternative views have been proposed, including the idea that increased cross-modal activations instead are simply due to an unmasking of existing inter-modal connections that does not necessarily come with any perceptual benefit, or serve to stabilize cortical dynamics in the absence of input signals (Singh et al., 2018). This is supported by the fact that reported perceptual benefits seem disproportionately small in relationship to the cortical area freed up by the missing sense, as well as by findings of worsened performance of sensory deprived individuals in some conditions (Bolognini et al., 2012; Kolarik et al., 2017; Kowalska & Szelag, 2006; Zwiers et al., 2001). As for olfactory sensory deprivation, (Peter et al., 2019) showed that individuals with congenital anosmia were better at detecting audio-visual temporal asynchronies, suggesting that the deleterious effects that olfactory deprivation has been shown to have on taste and trigeminal perceptual abilities (Frasnelli et al., 2010; Gagnon et al., 2014; Hummel et al., 2003) may be specific to the intrinsically connected chemical senses. Thus, our results of altered cross-modal activations and functional connectivity in individuals with CA may reflect an adaptive plasticity.

One possibility is that these cross-modal activations reflect the processing of object-related features in the piriform cortex. Compared to other sensory modalities, olfaction is arguably more dependent on information from other senses for object identification in that we are particularly bad at identifying odor sources without the help of relevant cues from our other senses (Desor & Beauchamp, 1974) (Cain, 1979; Zellner et al., 1991). In this view, the main task of the piriform cortex may be to process objects and to categorize input, with a preference for olfactory input but no dependence on concurrent or even previously experienced olfactory stimulation. The idea of a task-based brain organization where the purpose of the piriform cortex is first and foremost object processing, albeit with an olfactory preference, predicts strong cross-modal activations also in individuals born without a sense of smell. If the main task of the piriform cortex is to process object related information rather than olfactory input per se, its processing of sensory information from other modalities may be less dependent on olfactory experience. Our results are compatible with the idea of the piriform cortex as an object-processing region rather than a purely olfactory processing one, although we did not assess activation by non-object stimulation. Our stimuli consisted of sounds and pictures of objects, meaning that we did not test whether the piriform cortex would respond also to non-object stimulation such as for example noise. To firmly establish non-olfactory object processing in the piriform cortex, and to what extent it is preserved in congenital anosmia, future studies could compare to object versus non-object stimulation and for example assess decoding of object identity similarly as previously done for olfactory-linked stimuli (Kehl et al., 2024).

In summary, our results indicate that the cross-modal object processing in the piriform cortex previously demonstrated in normosmic individuals does not require any previous olfactory experience. In addition, we found that the congenitally olfactory deprived individuals had an altered functional connectivity during auditive and visual processing within the piriform cortex and between the piriform cortex and other brain regions. Considering that stronger cross-modal visual activation in the auditory cortex in hearing impaired individuals has been associated with poorer outcome after cochlear implant surgery (Paul et al., 2025), characterizing the cross-modal plasticity in anosmia has potential importance for future development of olfactory implants.

## ACKNOWLEDGEMENTS

Funding was provided by the Knut and Alice Wallenberg Foundation (KAW 2018.0152), the Swedish Research Council (2021-06527), and the European Union (ERC, D2Smell, 101118977), awarded to JNL. Data acquisition was supported by a grant to the Stockholm University Brain Imaging Centre (SU FV-5.1.2-1035-15). We thank Anja Winter and Katharina Prenner for their help with data collection.

## SUPPLEMENTARY MATERIAL

**Supplementary Table S1.** Full results from ROI analysis, per experimental group and ROI. Statistical comparison against baseline using Welch’s t-tests.

| Condition | ROI | Mean | SD | df | *t* | *p* | *d* |
| --- | --- | --- | --- | --- | --- | --- | --- |
| **CA** |  |  |  |  |  |  |  |
| *Visual* | aPir | .50 | .76 | 29 | 3.58 | .001 | .65 |
| | pPir | 1.21 | 1.04 | 29 | 6.39 | $5.5 \cdot 10^{-7}$ | 1.17 |
| *Auditory* | aPir | .73 | .99 | 29 | 4.03 | $3.6 \cdot 10^{-4}$ | 0.74 |
| | pPir | 1.19 | 1.29 | 29 | 5.05 | $2.2 \cdot 10^{-5}$ | 0.92 |
| **Control** |  |  |  |  |  |  |  |
| *Visual* | aPir | -0.22 | .87 | 29 | 1.35 | .186 | .25 |
| | pPir | .67 | .88 | 29 | 4.12 | $2.9 \cdot 10^{-4}$ | .75 |
| *Auditory* | aPir | .81 | 1.17 | 29 | 3.81 | $6.8 \cdot 10^{-4}$ | .69 |
| | pPir | 2.11 | 1.74 | 29 | 6.64 | $2.8 \cdot 10^{-7}$ | 1.21 |

CA - Congenital Anosmia; ROI – Region of Interest

